# Assessment of salivary microRNA by RT-qPCR: Challenges in data interpretation for clinical diagnosis

**DOI:** 10.1101/2024.02.06.579150

**Authors:** Marc Van Der Hofstadt, Anna Cardinal, Morgane Lepeltier, Jérémy Boulestreau, Alimata Ouedraogo, Malik Kahli, Pierre Champigneux, Laurence Molina, Franck Molina, Thi Nhu Ngoc Van

## Abstract

Salivary microRNAs (miRNAs) have been recently revealed as the next generation of non-invasive biomarkers for the diagnostics of diverse diseases. However, their short and highly homologous sequences make their quantification by RT-qPCR technique highly heterogeneous and study dependent, thus limiting their implementation for clinical applications. In this study, we evaluated the use of a commercial RT-qPCR kit for quantification of salivary miRNAs for clinical diagnostics.

**Methods:** Saliva was sampled from ten healthy volunteers for a time course analysis. A panel of six miRNA targets (with different sequence homologies) were analysed by one of the most commonly used commercially available RT-qPCR kit. Sensitivity and specificity of the tested miRNA assays were corroborated using synthetic miRNAs. The reliability of all tested assays to differentiate miRNA expression profiles were analysed, to statistically discriminate background noise from intrinsic individual signals.

**Results:** Significant variabilities in expression profiles of six miRNAs from ten healthy participants were revealed, yet the poor specificity of the assays offered insufficient performance to associate these differences to biological context. Indeed, as the limit of quantification (LOQ) concentrations are from 2-4 logs higher than that of the limit of detection (LOD), the majority of the analysis for salivary miRNAs felt outside the quantification region. Most importantly, a remarkable number of crosstalk reactions exhibiting considerable OFF target signal intensities was detected, indicating their poor specificity and limited reliability. However, the spike-in of synthetic miRNA increased the capacity to discriminate endogenous salivary miRNA at the LOQ concentrations from those that were significantly lower.

**Conclusions:** Our results demonstrate that comparative analyses for salivary miRNA expression profiles by this commercial RT-qPCR kit are most likely associated to technical limitations rather than to biological differences. In particular, assessment of fundamental parameters including LOD, LOQ and crosstalk of each assay is strictly necessary to interpret observed variations. The standardization of rigorous sample handling and experimental design according to technical parameters of each assay plays a crucial role in reducing data inconsistencies across studies. However, further technological breakthroughs are still required to overcome discrepancies in order to accelerate the translation of salivary miRNAs for clinical applications.

## Introduction

In the last few decades, human saliva has attracted a great attention in the medical diagnostic field as a bio-fluid that provides access to various biomarkers in a non-invasive manner. Similar to peripheral blood, saliva contains a high versatility of circulating molecules that allows early diagnosis of systemic disorders [1,2]. For clinical practice, saliva offers a number of advantages over blood sampling such as a remarkable stability, simple sample collection, possibility for auto-sampling, and practical transportation and storage condition [1]. In addition, saliva has a high turnover rate exhibiting fast bidirectional exchange of biomarkers, which opens access to recent biological events. However, the high turnover rate is at the cost of harbouring large biochemical and physical dynamics among samples [2] and the presence of buccal mucosa and non-human derived (*i*.*e*. oral bacteria) biomaterial that mask the low concentration of circulating biomarkers. Altogether, this makes the quantification of salivary biomarkers highly challenging, requiring exceedingly robust assays [3]. Nevertheless, the concentration levels of salivary biomarkers of different natures including DNA, miRNA, RNA, protein and metabolite have been identified to be associated to a broad range of diseases [4-7] via numerous clinical studies, demonstrating the significant clinical interest of precisely quantifying these biomarkers.

Recently, salivary miRNAs have been considered as the next generation of non-invasive biomarkers for the diagnostics of diverse diseases [8,9]. They are small non-coding RNAs that regulate gene expression at the post-transcriptional level, playing a crucial role in every fundamental aspect of cellular function [10]. The relationships of their regulation with the disease onsets [8,9] and their presence in the extracellular circulation [11-13] make them available for various liquid biopsies enabling non-invasive medical assessment [14,15]. For instance, signatures of salivary miRNAs have been recently associated with the diagnosis of mild traumatic brain injury (mTBI) [6,16-18], cancers [12,19,20], endometriosis [21,22], neurodegenerative [23], metabolic [24] or systemic diseases [7] along with many others [25-29]. However, due to their small size (19-25 nucleotides), their high sequence similarity and their complex gene regulation process [10], the correlation between their expression levels and disease stages is often non-linear, causing it an issue of constant debate [8,15,30]. The majority of the miRNA studies have been standardized by microarray or Next Generation of Sequencing (NGS) methods followed by subsequent validation using various customized RT-qPCR approaches [31-36]. Notwithstanding the practicality, rapidity and cost effectiveness [8] of the RT-qPCR technique, its application in the quantification of miRNAs faces methodological inconsistencies in both detection and data normalization [37-38], making data interpretation controversial [39] and study dependent [18]. These unsettled topics are the main issues that restrains the use of miRNAs in current clinical practice.

Understanding technical issues of a given assay for quantification of salivary miRNA to navigate corresponding solutions would accelerate the clinical translation process for salivary miRNA biomarkers. To this aim, in this study we evaluated the use of a wildly used commercial RT-qPCR kit [31] for quantification of salivary miRNAs by determining its capacity to specifically discriminate miRNAs with various homologous degrees. A panel of six miRNAs, namely hsa-Let-7a-5p, hsa-Let-7f-5p, has-mir-148a-3p, has-miR-26b-5p, hsa-miR-107 and hsa-miR-103a-3p was chosen based on both their clinical values [16,40,41] and their sequence homology. RT-qPCR assays were performed on both miRNA targets extracted from saliva of healthy volunteers and synthetic ones. Fundamental technical parameters such as sensitivity, specificity, limit of detection (LOD), limit of quantification (LOQ) and cross-reactions of all assays were characterized and included in the data interpretation process.

## Materials and Methods

### Study approval and saliva collection

Ten healthy males with ages ranging from 18 to 40 were recruited in the Sys2diag laboratory based in Montpellier, France, according to the personal protection committee (CPP) with registered number 23.00930.000169 (NCT06149351 on www.clinicaltrials.gov). All subjects were informed, signed and consented in accordance with the CPP prior to the recruitment. Saliva collection and analyses were performed with approved protocols. Participants were asked to refrain from eating, drinking or smoking for at least 30 min prior to saliva collection. During the 3 months of the study, a total of four collections (one collection every 2-3 weeks) were performed at the same time of the day. Approximately 2 mL of unstimulated saliva was collected from each participant and stored at 4°C for a maximum of 2 hours prior small RNA extraction. The appearance of each individual saliva was visually inspected while its viscosity was estimated via its flow resistance using a combitip®.

### Salivary miRNA extraction and quantification

Total small RNA was extracted from 250 μL of whole saliva using miRNeasy Serum/Plasma Kit (Qiagen) according to the manufacture instructions (except for the elution step, where the extracted salivary small RNA was collected in 20 μL nuclease free water). In brief, 250 μL of whole saliva from each participant was homogenized in 1 mL of Qiazol solution, and was followed by the addition of 5.6 × 10^8^ copies of UniSP6 miRNA as a technical control for the extraction process, unless mentioned otherwise. Chloroform purification was followed by RNA precipitation by isopropanol, which was then loaded into a miRNeasy column, where only small RNA fragments (<200 bp fragments) are retained following multiple washes.

Total small RNA concentration was quantified by Nanodrop One (Thermo Scientific, Wilmington USA). Small fragments of eluted RNA was confirmed by Labchip using small RNA assay (PerkinElmer).

### Synthetic miRNA Targets

Six miRNAs, including hsa-Let7a-5p, hsa-Let7f-5p, hsa-miR-148a-3p, hsa-miR-26b-5p, hsa-miR-107 and hsa-miR-103a-3p, were selected according to their sequence similarity and clinical relevance. All the six synthetic miRNAs were purchased from Integrated DNA technologies (IDT, Europe). All sequences and annotations are available in the supplementary data section (Table S1).

### Reverse transcription

miRCURY LNA RT Kit employing poly (A) polymerase for tailing RNA prior to an universal reverse transcription (RT) using poly T primer was purchased from Qiagen. 10 μL RT reactions were prepared in 96 well plates containing 2 μL of 5X reaction buffer, 1 μL 10X reverse transcriptase and 1 μL of synthetic or extracted small RNA at desired concentrations. The reactions were incubated in a peqSTAR 96X thermocycler (Ozyme, Montigny-le-Bretonneux France) at 42°C for 60 min, followed by a denaturation step at 85°C for 5 min and stored at - 20°C until used. No template and no reverse transcriptase enzyme negative controls were included in each run. UniSp6 miRNA was included when needed as plate calibrator. All samples and control conditions were run in duplicate.

For the analysis using 50 ng of extracted small RNA as input, all extracted samples were normalized to 50 ng/μL and then 1 μL was added to the RT-qPCR reaction, except for P5 and P6 samples whose concentration were inferior to 50 ng/μL, and hence higher volumes were required.

### Q-PCR quantification and analysis

miRCURY LNA SYBR Green PCR Kit and target specific primers (miRCURY LNA miRNA PCR assays) were purchased from Qiagen. 10 μl qPCR reactions were prepared in 384 multi-well plates containing 5 μL of qPCR master mix; 1 μL of corresponding primers (miRCURY LNA miRNA PCR Assay) and 3 μL of 10X diluted RT product. QPCR negative controls (No-template reactions) were included for each assay. All samples and control conditions were run in duplicate. Real-time qPCR thermal cycling reactions were performed by the LightCycler 480 (Roche, Meylan France) directed by the LightCycler 480 Software (version 1.5.1.62). Thermal cycling conditions were: Pre-Incubation for 2 minutes at 95°C, 40 cycles of amplification (95°C for 10s, 56°C for 60s). Ct values were analysed by Abs Quant/2nd Derivative Max of the same software. Ct values of UniSP6 was verified prior to all analysis when plate calibrator was needed. All Ct value at 35 was considered as noise.

### Statistical Analysis

Scipy version 1.11.2 was used on python 3.10.4 to do statistical analysis. All statistical test were done using Mann-Whitney U (scipy.stats.mannwhitneyu) with asymptotic method (*i*.*e*. p-value calculated by comparing to normal distribution and hence correcting for ties).

## Results

### Saliva physical characteristics and their small RNA content do not affect RNA extraction efficiencies

Following each sampling, physical characteristics of individual saliva samples were visually inspected prior to small RNA extraction. Results revealed that saliva from a given participant (P) had a typical physical property harbouring a given level of cesia and viscosity, which remained unchanged throughout the 4 samplings of the study (Figure 1A&B). Concentrations of total extracted small RNA varied both among participants and sampling times (Figure 1C), although no significant difference among the average concentrations of the four samplings was observed (Figure S1). We highlighted consistently low concentrations across the study for P5 and P6 samples (Figure S2), whose saliva were both liquid and transparent. Efficiencies for every individual small RNA extraction were technically controlled by quantifying the retained amount of the UniSP6 miRNA that had been equally spiked-in preceding the extraction process. Consistent RT-qPCR results (Ct = 21.7 ± 0.07) using 1μL of extracted small RNA from all samples revealed a comparable extraction efficiency for all participants (Figure 1D), which was conserved throughout this study (Figure S3). More precisely, upon standardizing the total RNA concentration with respect to 1 μL condition, UniSP6 detection remains linear demonstrating that the small RNA content of a given sample does not affect its extraction efficiency (Figure S4). Subsequently, we validated that the addition of this spiked-in miRNA had no influence on the extraction efficiency for the six target miRNAs. To this aim, we chose three samples (P2, P6 and P7) whose small RNA contents are in different range (Figure 1C) to perform the RNA extraction procedure in the presence or absence of the spiking UniSP6 miRNA (Figure S5). Results of RT-qPCR analysis using 50ng of each extracted small RNA from these samples showed that spike-in UniSP6 had no differential effect on the six assessed miRNAs as the observed Ct values increased by only less than 1% (ΔCt 0.2 ± 0.09), and (Figure S5C).

**Figure 1.**
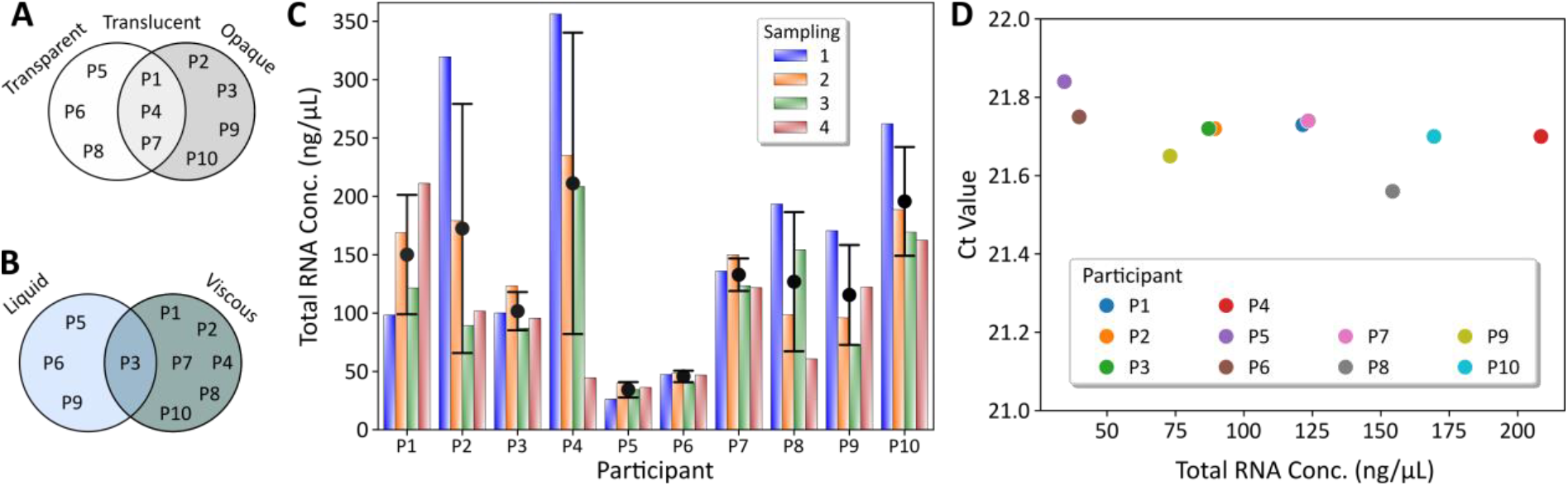
Physical properties and small RNA extraction of saliva samples from 10 healthy participants. (**A**) Cesia and (**B**) viscosity of the saliva samples prior RNA extraction. (**C**) Total small RNA extracted from 250 μL of saliva for each participant and for the 4 sampling points. Error bars show one standard deviation. (**D**) Detection of spiked artificial UniSP6 miRNA prior to RNA extraction for sampling three. 1 μL of each extracted RNA sample was used as input. P1-10: Saliva sample from participants 1 to 10.

### Salivary miRNA expression profiles vary across samplings and participants

A panel of six miRNAs was analysed on each saliva sample (10 participants and 4 temporal sampling points, n=40). 50 ng of extracted small RNA was used as input for universal poly(A)-tailed RT reactions followed by target specific qPCR quantification (Qiagen). Figure 2A and Table S2 show the heterogeneous expression profiles obtained from detection signals of the six target miRNAs with averaged Ct values for the four samplings ranged from 22 to 29 (±1.5), which are within the acceptable detection ranges considered by the community^19,42^. However, the six miRNA targets shared a similar detection pattern among the four sampling points (Figure 2B). We observed a reduction in Ct value from the sampling 1 to sampling 3, followed by a restoration in the sampling 4 whose Ct values fell between the first two samplings. Even if we had previously observed no significant difference between the average of the extracted total RNA concentrations across temporal sampling (Figure S1), Ct values on 50 ng samples revealed the contrary, showing a significant difference between sampling 1 and 3 for all miRNAs, and between sampling 3 and 4 for half of the miRNA assessed, namely hsa-Let-7a-5p, hsa-miR148a-3p and hsa-miR103a-3p (Figure S6A). Statistical analysis for individual miRNA targets revealed the apparition of two major groups, a lower Ct value group containing hsa-Let-7a-5p and hsa-Let-7f-5p, and a greater Ct value group containing hsa-miR148a-3p and hsa-miR107 (Figure 2C and S6B). In between both groups, we found hsa-miR26b-5p and hsa-miR103-3p, which shared proximity with the lowest and the greatest Ct value group, respectively. However, the evaluation of the Ct values with respect to individual participants showed that the average miRNA signals (Figure 2B) partially masked the ones of individual participants (Figure 2C). Further analysis on these miRNA expression profiles revealed significant differences among participants, which can be clustered into three major groups: A) high Ct values (P5 and P6), B) moderate Ct values (P8 and P9) and C) low Ct values (P1, P2, P3 and P4), with participant P7 and participant P10 between the later groups, being adjacent to group B&C and group C, respectively (Figure S7A). We noted that Ct values of these groups were in close relation to the dilution factor required to achieve the 50 ng of extracted small RNA for RT-qPCR experiments (Table S3). More precisely, the first group presented small dilution factors (7x and 9x), the second group intermediate (25x and 23x, followed by P7 with 26.5x) and the third group highest (ranging from 30x to 42x, with the exception of the P3).

**Figure 2.**
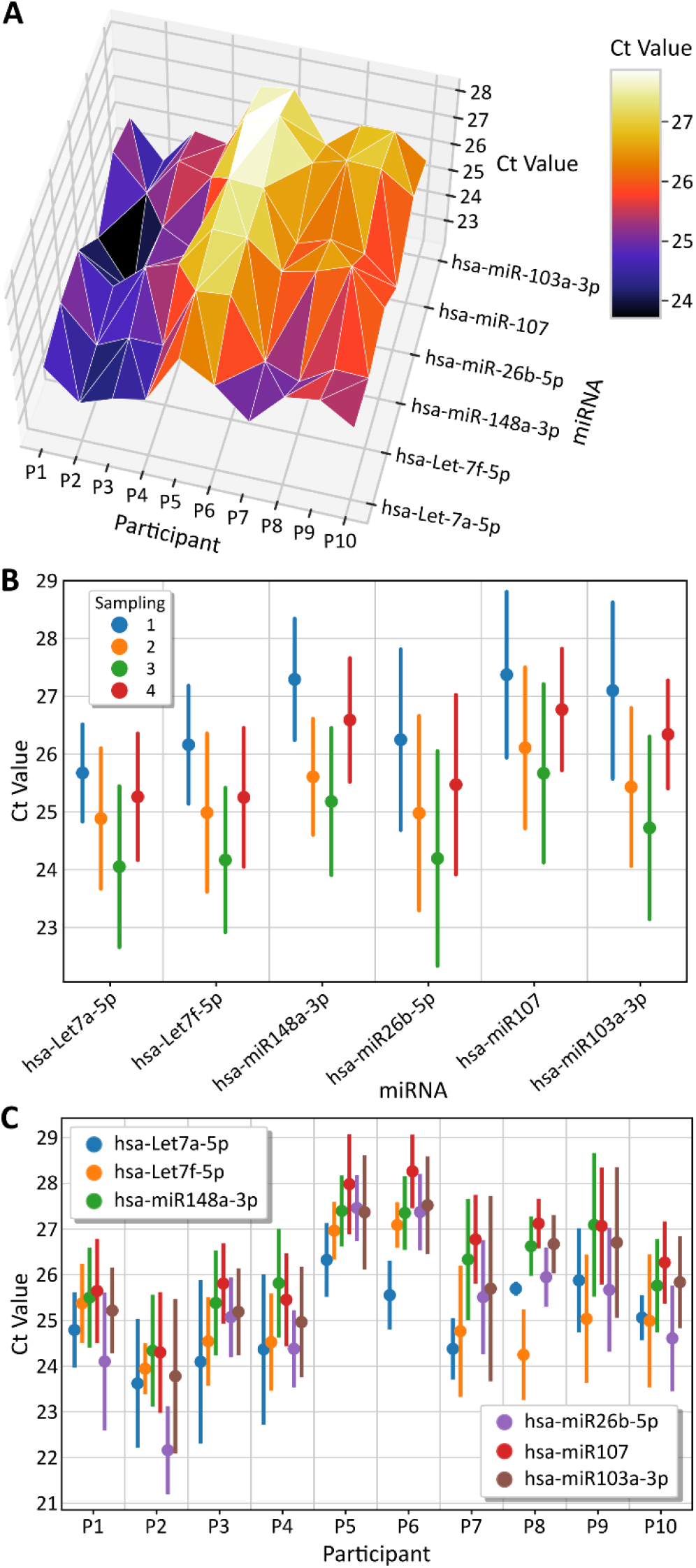
RT-qPCR shows high variability of salivary miRNA expression profiles within 10 healthy participants. **(A)** Average Ct values of the four sampling points through which the six miRNAs was assessed on 10 participants. Expression profiles of the six analysed miRNAs across **(B)** time and **(C)** different participants. All analysis was perform with 50 ng of total extracted small RNA. Error bars show one standard deviation.

To fully evaluate the effect of the dilution factors on miRNA detection signals, we next performed dose dependent response experiments on three samples (P2, P6, P7). The six miRNA assays were performed using different extracted small RNA input concentrations for RT-qPCR, ranging from 1 ng up to 300 ng. Results demonstrated linear behaviour for all the six miRNAs for three participants (Figure S8). However, while participants P2 and P7 showed high correlations coefficients (R^2^>0.99, except for hsa-Let-7f-5p in P7), participant P6 had lower coefficients, ranging from 0.94 to 0.99. Particularly, at the highest concentration (300ng), Ct values were lower than predicted by the linear fit (Figure S8B). Similarly, although the six tested qPCR assays had comparable efficiencies, overall higher efficiencies were observed for participants P2 and P7 compared to participant P6 (Figure S9). Nevertheless, when averaging the three participants, we observed that miRNA expression profile was conserved throughout the tested concentration range (from 1 ng up to 300 ng of extracted small RNA) (Figure S8D). Indeed, the detection of both UniSP6 (Figure S4) and the six miRNA targets (Figure S10) detected linearly at different working dilutions.

### High cross-reactivity limits the reliability of the miRNA assays

To determine to what extent the variability observed in endogenous salivary miRNA quantification was associated to biological factors, we investigated the reliability of the assessed assays. To this aim, we investigated the sensitivity and specificity of the six miRNA assays using their corresponding synthetic targets. The limit of detection (LOD) for each miRNA assay was determined by signals obtained at the lowest target concentration in a serial dilution of concentration ranging from 1 to 10^12^ copies/μL (Figure 3A). Results demonstrated very high sensitivities, allowing to detect down to at least 1 copies/μL of miRNA target from their negative controls (No detection or Ct Value = 35). However, we noted that under a given concentration, the detection signals was no longer linear, and hence we defined it as the limit of quantification (LOQ). The LOQ concentrations for hsa-Let-7a-5p, hsa-miR-148a-3p, hsa-miR26b-5p and hsa-miR-107 assays were higher (10^5^ copies/μL) compared to that of hsa-Let-7f-5p and hsa-miR-103a-3p assays, which decreased down to 10^4^ and 10^2^ copies/μL, respectively. Calculation of the RT-qPCR efficiencies between the LOQ and the Ct saturation point (*i*.*e*. Ct value = 5) revealed high efficiencies for all miRNA assays, ranging from 98.5 and 112.5% (Figure S11 & S12), which fall within reasonable limits for PCR exponential amplification.

**Figure 3.**
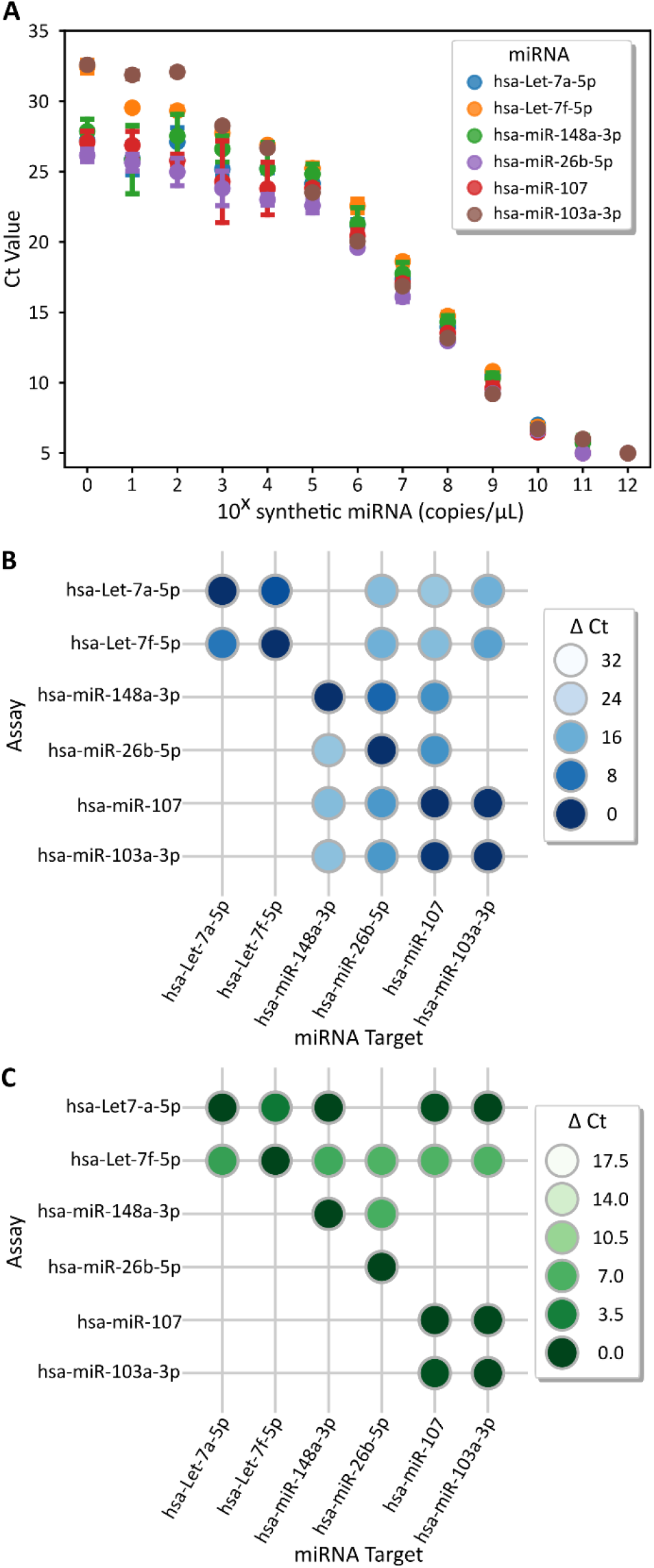
Sensitivity and specificity limitations of the six miRNA assays. (**A**) Serial dilution from 1 to 10^12^ copies/μL with synthetic miRNA target for each miRNA assay to determine their limit of detection (LOD) and their limit of quantification (LOQ). Cross reactions between miRNA assays and synthetic targets at (**B**) 10^9^ copies/μL and (**C**) 10^5^ copies/μL. ΔCt values are calculate from Figure S13. Empty spaces represent no crosstalk (no detection or Ct Value = 35). Data obtained from a duplicate experiment. ΔCt values = Ct^OFF target^ – Ct^ON target^.

Due to the short size, and hence high sequence similarity [15], miRNA detection is prone to crosstalk. To undercover the potential crosstalk among assays and targets used in this study, we compared the detection signal of each assay reporting on its corresponding miRNA target (ON target) to that of the detection on the other targets (OFF target) of the panel. ΔCt values were calculated between the Ct value of the OFF target and the one ON target as indicator of their cross-reactivity (crosstalk). Firstly, we assessed the crosstalk contribution at 10^9^ copies/μL, a concentration that corresponds to the middle of the quantification zone (Figure 3B & S13A). Out of the 30 possible crosstalk combinations, only 12 of them showed no crosstalk (No detection or Ct Value = 35) while the rest (60%) presented significant crosstalk. The most striking crosstalk was observed between hsa-miR-107 and hsa-miR-103a-3p assays with respective ΔCt of 0.21 and 0.84, making their discrimination impossible. In the same line, it is very unlikely to discriminate hsa-Let7a-5p from hsa-Let7f-5p as these assays also presented high crosstalk exhibiting ΔCt values of 3.9 and 8.7, respectively. The rest of crosstalk combinations presented variable ΔCt, ranging from 6.5 up to 25.4. We next assessed crosstalk contribution at 10^5^ copies/μL, which corresponds to the LOQ concentration for four out of the six miRNA assays. At this concentration, although the number of crosstalk reduced down to 40%, most of the retained OFF targets drastically increased their significance (the average ΔCt decreased from 13.14 down to 3.6) (Figure 3C & S13B). As for the two pairs: hsa-miR-107 vs hsa-miR-103a-3p, and hsa-Let7a-5p vs hsa-Let7f-5p, their maximal crosstalk at 10^9^ copies/μL remains unchanged when concentration was lower to 10^5^ copies/μL.

We then further verified the respective crosstalk throughout the whole serial dilution (concentration ranging from 1 to 10^12^ copies/μL) for the two assays hsa-Let7a-5p and hsa-Let7f-5p. As shown in Figure 4, crosstalk remained generally stable throughout different concentrations with an average ΔCt value of 3.78 ±1.02 for hsa-Let7a-5p (Figure 4A) and 10.29 ± 1.20 for hsa-Let7f-5p (Figure 4B). However, we noticed two tendencies: first, when the concentration of the OFF target decreased the error value increased; and second, a transition at the LOQ, where lower concentrations presented erratic responses. Re-calculation of the average ΔCt within the limit of the quantification zone didn’t considerably change, being 3.93 ± 1.01 for hsa-Let7a-5p and 10.02 ± 1.1 and for hsa-Let7f-5p.

**Figure 4.**
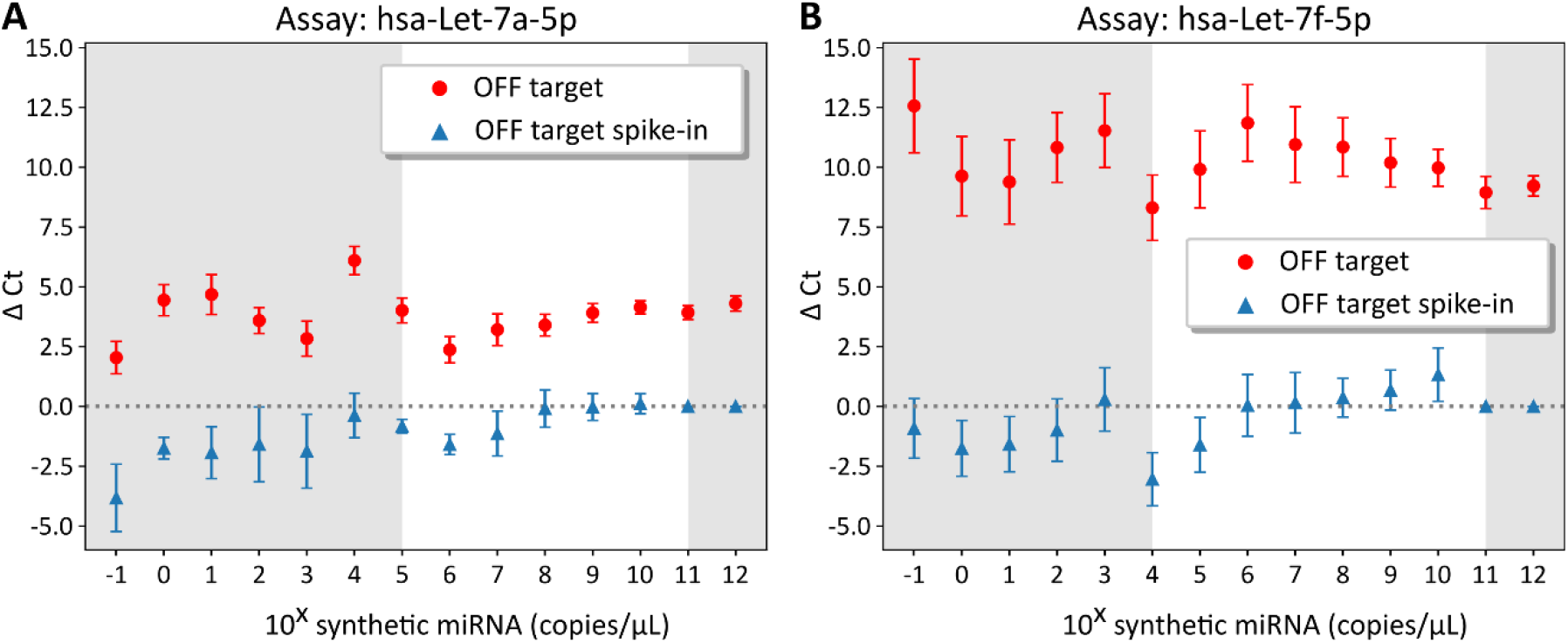
Conserved crosstalk for hsa-Let-7a-5p and hsa-Let-7f-5p within the quantification region. ΔCt values obtained when comparing a serial dilution from 10^−1^ to 10^12^ copies/μL of the corresponding synthetic miRNA (ON target) with its cross reaction (OFF target, red circles) for (**A**) hsa-Let-7a-5p assay and (**B**) hsa-Let-7f-5p assay. The ΔCt by the addition of 10^7^ copies/μL of OFF target on the serial dilution of ON target was also calculated (blue triangles). Error bars show one standard deviation from a duplicate experiment. Red circles = Ct^OFF target^ – Ct^ON target^, blue triangles = Ct^ON target + OFF target at 10^7 copies/μL^ – Ct^ON target^. Grey region delimits the quantification region.

Knowing the OFF target effect of both assays, we decided to evaluate the LOD of each assay in the presence of the OFF target at the LOQ concentration. To do so, we re-assessed the serial dilution of the ON target in the presence of 10^7^ copies/μL of the OFF target (Figure 4). Results revealed that the presence of 10^7^ copies/μL OFF target did not affect the detection of the ON target, for both hsa-Let-7a-5p and hsa-Let-7f-5p assays. For the hsa-Let7a-5p assay, the greatest crosstalk influence of the OFF target was observed at 10^6^ copies/μL of the ON target (1 log difference), and a partial crosstalk at its adjacent concentrations (*i*.*e*. 10^5^ and 10^7^ copies/μL). Similarly, for hsa-Let7f-5p assay, the greatest crosstalk contribution was observed at 10^4^ copies/μL of the ON target (3 log difference), showing a greater resilience of hsa-Let7f-5p assay to crosstalk compared to the hsa-Let7a-5p assay. These results are in agreement with the average ΔCt value of 3.9 (⌼ 1 log) and 10.0 (⌼ 3 logs) for the OFF target of the hsa-Let7a-5p and hsa-Let7f-5p assays, respectively.

### Only a minority of the miRNA detections falls within the quantification regime

Up to this point, we reconsidered the term “good range of detection” stated previously (Figure 2) based on the linear quantification zones for the six miRNA assays (Figure S11). Indeed, upon taking into consideration the LOQ, we observed that out of the 240 quantified samples (from the total of 10 participants, 4 samplings, 6 miRNA assays), 42% of them were analysed outside the quantification zone (*i*.*e*. Ct values higher than the LOQ), which included all the 40 samples analysed with the hsa-miR-107 assay. Subsequently, we also noticed that 29% of the results were at the respective LOQ of the miRNA assay, leaving assays such as hsa-Let-7a-5p and hsa-miR-148a-3p analysed either outside or at the LOQ. Yet, of the 29% obtained Ct values within the linear quantification zone, the hsa-miR103 assay was the sole to be completely analysed within this zone (as a result of its lower LOQ). These results indicated that endogenous salivary hsa-miR103-3p was present at concentrations between 3.8Ð10^3^ and 5.4Ð10^4^ copies/μL, and that only participant P6 showed partial significant differences with respect to the other participants (Figure S7B).

Given the high Ct values obtained for endogenous salivary hsa-Let7a-5p analysis, we decided to spike-in the synthetic hsa-Let7a-5p to surpass the LOQ concentration. To this end, we chose 4 spiking concentrations; two that were significantly under the LOQ (10^−1^ copies/μL and 10^2^ copies/μL), one at the LOQ (10^5^ copies/μL) and the last one at 10-fold higher than the LOQ (10^6^ copies/μL). As expected, the higher the concentration of spiked-in synthetic hsa-Let7a-5p miRNA, the higher the ΔCt value with respect to the endogenous salivary sample (Figure 5a & S14). However, two distinctive groups appeared; a group (P1 and P3) whose detection signals were not altered until upon the addition of 10^5^ copies or above of the synthetic target and a second group (P4, P5, P6, P8 and P9) that showed alteration in detection signal directly with the addition of only 10^−1^ copies/μL (*i*.*e*. 1 copy in the reaction) of the synthetic target. To understand these shifts, we calculated the ΔCt value between the spiked-in salivary samples and the corresponding synthetic serial dilution (Figure 5B). Results showed that only for participants P1 and P3 the addition of 10^6^ copies/μL had a negative effect on the detection of hsa-Let7a-5p compared to the synthetic target, while for the remaining participants, this effect was observed at 10^5^ copies/μL.

**Figure 5.**
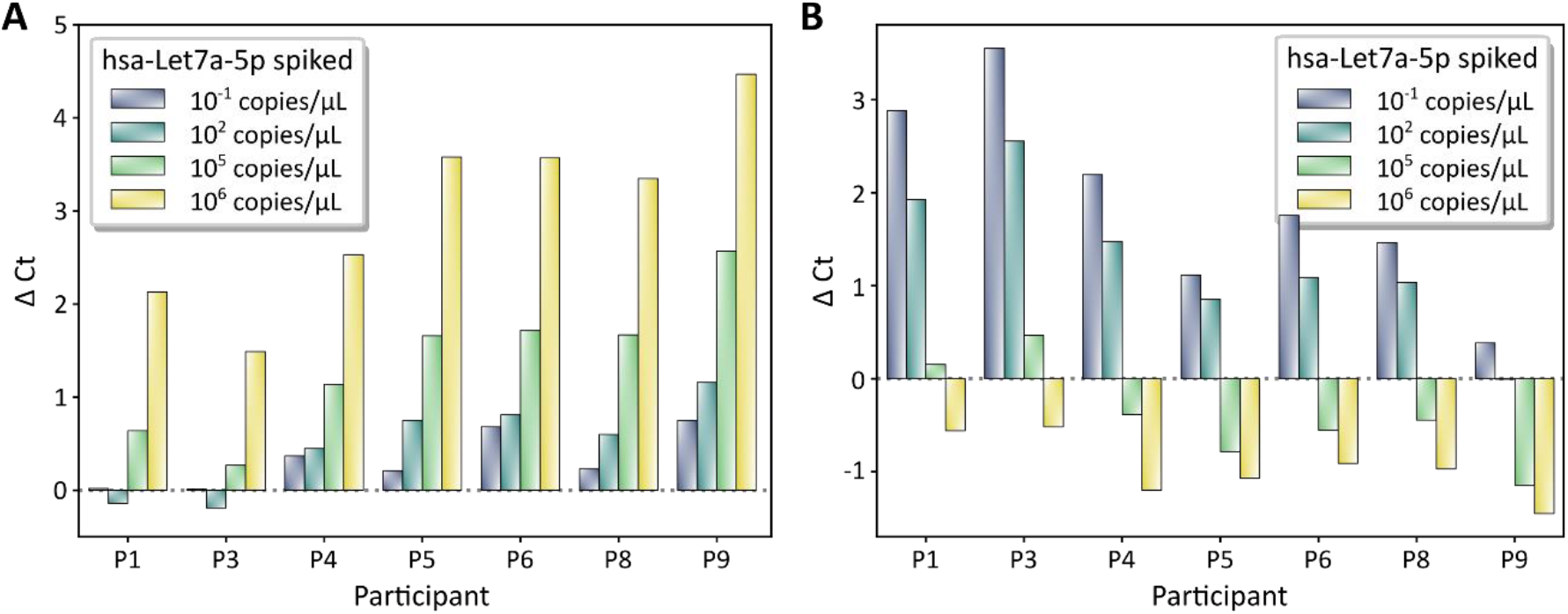
The spiking of synthetic hsa-Let-7a-5p miRNA allows semi-quantification at LOQ concentrations. (**A**) ΔCt values of spike-in synthetic hsa-Let-7a-5p miRNA in 50 ng RNA extract with respect to the non-spike saliva sample in a range of spike-in concentrations. ΔCt were calculated as Ct^Endogenous^ - Ct^Endogenous + synthetic^. (**B**) ΔCt values of spike-in synthetic hsa-Let-7a-5p miRNA in 50 ng RNA extract with respect to same synthetic miRNA concentration in the absence of sample. ΔCt were calculated as Ct^synthetic^ - Ct^endogenous + synthetic^. Data determined from Figure S14.

## Discussion

Salivary miRNA biomarkers have become recently relevant to the medical diagnostic community due to their easiness in sample collection and their expression signatures in diverse diseases [4,6,7]. However, the lack of standardized approaches in experimental design, method of analysis and data interpretation coupled with inter-individual variability has led to the generation of considerable study-dependent and controversial data [37,39]. We evaluated the capacity to detect salivary miRNAs of a commercially available kit, which is widely used in both research and clinical studies [16,19,40]. We observed significant variability between miRNA targets and participants, although these variations cannot be fully attributed to biological fluctuations, reinforcing the need for an accurate data interpretation in clinical studies.

Our data shows that although saliva obtained from different participants vary in physical properties and small RNA concentration, this heterogeneity did not affect the efficiency of the small RNA extraction process. Interestingly, we did observe that transparent and liquid samples presented smaller variations in small RNA content throughout the four samplings and significantly lower small RNA concentrations (Figure 1C & S2). However, this low concentration of extracted small RNA did influence the downstream RT-qPCR analysis of the sample. Indeed, serial dilutions of total small RNA extracted from participant P6 also presented slightly lower correlations coefficients (Figure S8) and partially lower efficiencies (Figure S9) for RT-qPCR. Since higher deviations are observed at high concentrations of small RNA (300 ng), we hypothesize that the use of higher volumes (⌼4-fold greater compared to participant P2) would carry on more chemical residues from the purification process, which may affect the efficiency of the RT-qPCR. For this reason, and due to the conservation of the miRNA detection profile at different small RNA concentrations (Figure S8D), we propose 50 ng as input concentration for RT-qPCR as a good compromise when using human saliva samples.

Relying on already established potential miRNA biomarkers for the diagnostic of brain concussion [6,16-18], we quantified a representative panel of six miRNA (hsa-Let-7a-5p, hsa-Let-7f-5p, has-mir-148a-3p, has-miR-26b-5p, hsa-miR-103a-3p and hsa-miR-107) in saliva from ten participants at four sampling points. Interestingly, we firstly noted that although no significant difference among the average concentrations of the extracted small RNA was observed throughout the four samplings of the study (Figure S1), upon standardization to 50 ng, a significant fluctuation of miRNA detection signal across time was obtained (Figure 2B). This could be accounted to the modification in the presence of other small RNAs (including those are not analysed in this study), indicating that a direct relation between total extracted small RNA concentration and miRNA concentration cannot be done. Secondly, we also observed that the temporal tendency is considerably marked for the six miRNAs assessed. We initially hypothesized that this variation may have been attributed to an environmental effect that had homogenously affected all participants throughout the three months of the study. However, given the technical limitations described in the subsequent sections, there was not sufficient robustness to validate as the unique hypothesis.

Statistical analysis of the six miRNA targets significantly clustered them into two major groups: a low Ct value (hsa-Let-7a-5p and hsa-Let-7f-5p) and high Ct value (hsa-miR148a-3p, and hsa-miR107) one. Yet, as expected, this average miRNA expression profile partially masked the one of individual participant (Figure 2C), demonstrating the need for individualized data analysis and the risk in data clustering, especially when defining a molecular profile in clinical diagnostics. Similarly, we observed statistically significant differences among participants that have been related to their dilution factor (Table S3), and hence to their extracted small RNA concentrations. In particular, that the two participants P5 and P6 stand out as an individual group (Figure S7) with high Ct values. Since miRNA expression profiles of P5 and P6 were very similar to the rest of the participants (Figure 3C), there was no clear difference between the high efficiency miRNA assays and the lower efficiency ones analysing P6 sample (Figure S9) especially at high input concentrations (300ng), where assay efficiencies are mostly affected. Therefore, we do not discard the higher Ct values of P5 and P6 due to technical limitations when using 50 ng as input. Even if miRNA expression profiles were independent from the input concentrations of extracted small RNA (Figure S8D), samples with low small RNA concentrations face a greater technical risk of being outliers and therefore should be critically taken into consideration for data handling. Given the high heterogeneity in extracted small RNA concentrations (up to 13-fold difference) and the RT-qPCR impairments due to either excessive or insufficient small RNA concentration, we discourage the use of a fixed volume of small RNA for RT-qPCR as a normalizing method.

Given the particularly small size and the possibly low abundance of the circulating miRNA targets, the fundamental parameters of the employed RT-qPCR assays should be carefully investigated prior to drawing conclusions. Our results revealed that the LOQ concentrations for the six miRNA assays were significantly higher compared to that of their LOD ones (Table S4). Consequently, comparison of LOQ values to Ct values obtained from saliva samples revealed that only 40% of the obtained results were within the quantitative region. In particular, we could only infer quantitative information from hsa-miR-103a-3p assay, which demonstrated 1.4-fold difference between highest and lowest extracted small RNA concentration. These results highlight the technical limitations of the assays, which subsequently reduce the quantitative reliability of the obtained Ct values due to the insufficient sensitivity. Although small RNA concentration used could have been increased up to 200 ng (4-folds compared to 50 ng) to decrease Ct values and hypothetically increase up to 70% the number of results within the quantitative region, 4 out of the 6 miRNA assays would still not have all the values within the quantitative region, in addition to the cost of reducing qPCR efficiency as RNA input concentration increases.

In addition to sensitivity issues, RT-qPCR assays are commonly known to suffer from poor specificity for short targets such as miRNA [43]. Our results clearly demonstrated this phenomenon, where the cross-assessment of the six miRNA assays revealed considerable cross-detection, ranging from 60% at 10^9^ copies/μL (a concentration in the middle of the quantification zone, Figure 3B) down to 40% at 10^5^ copies/μL (the LOQ concentration for 4 out of 6 assays, Figure 3C). However, we observed that this reduction in cross-detection was at the cost of significantly increasing the significance of the remaining ones (*i*.*e*. worse specificity). In particular, and as expected, we observed that miRNA assays for targets whose sequences differed by only one nucleotide (*i*.*e* hsa-Let-7a-5p from hsa-Let-7f-5p and hsa-miR107 from hsa-miR-103a-3p) were unable of discriminating one from each other (Figure 3C). In particular, we observed that for the pair of hsa-Let-7a-5p and hsa-Let-7f-5p, crosstalk remained relatively constant throughout the serial dilution of the OFF target from 1 to 10^12^ copies/μL (Figure 4). Interestingly, we also noticed that the error was increasing as OFF target was decreasing, until the LOQ where the experimental errors stabilized. We attribute this behaviour to the stochastic (*i*.*e*. non-specific) nature of the enzymes [44] and the priming system of the assay, where at concentrations sufficiently low, the stochastic behaviour predominates over the deterministic behaviour, decreasing the robustness and linearity of amplification techniques [45,46]. Nevertheless, this high cross-detection, when carefully assessed, could beneficially be used for data interpretation. For example, although hsa-miR107 can only be quantified down to 10^5^ copies/μL with the hsa-miR107 assay, due to the high crosstalk of hsa-miR107 on hsa-miR-103a-3p assay, we can use the hsa-miR-103a-3p assay to reject the presence of hsa-miR107 at lower concentrations (as low as 3.8Ð10^3^, the LOQ of hsa-miR-103a-3p). Note that this statement allows us to decrease the capacity of rejecting the presence of hsa-miR107 by at least one order of magnitude only if hsa-miR-103a-3p is also absent.

To favour the ON target amplification at the LOQ without introducing more complexity to the matrix, hence increasing signal to noise ratio, we propose to spike-in synthetic miRNA. In particular, by spiking synthetic hsa-Let7a-5p in the extracted small RNA samples we intend to acquire semi-quantitative information, which would allow us to discriminate samples that were on the LOQ from those that were significantly lower. The rationality behind this statement is that the closer (or higher) the endogenous miRNA concentration is compared to the added synthetic miRNA, the lower the impact the added synthetic miRNA will have on the Ct value (*i*.*e* lower ΔCt). For example, the addition of 10^6^ synthetic copies/μL to a sample with endogenous miRNA at 10^4^ copies/μL (ratio 100:1) will largely affect its Ct value. However, as the endogenous miRNA concentration increases to 10:1 and 1:1 ratios (endogenous miRNA at 10^5^ copies/μL and 10^6^ copies/μL, respectively), the lower the variation in Ct value would be observed, since the total concentration of miRNA (endogenous plus synthetic) does not increase as significantly. With this rationality, we hypothesize that participant P1 and P3 have a concentration ⌼10^5^ copies/μL (although there is 0.68 ΔCt between them) while the other participants (P4, P5, P6, P8, P9) have concentrations < 10^5^ copies/μL, since the addition of 10^6^ synthetic copies/μL had low effect on the CT values of participant P1 and P3 but a significant effect for the other participants (Figure 5B). We do not venture in predicting the concentration of the latter group (or their order) since the addition of 10^5^ synthetic copies/μL is at the LOQ and hence is highly prone to error.

Taken together, our results demonstrate the importance to address the reliability of the chosen RT-qPCR kit when quantifying salivary miRNAs, since data can be highly mislead by technical limitations. Including the technical constraints, such as input concentration ranges, LOD, LOQ, PCR efficiency and cross-detection (and others within the MIQE guideline [47]), are necessary to support data interpretation, allowing to distinguish the biological relevance of the RT-qPCR results from variations associated with technical and methodological limitations.

## Conclusions

In this work we have shown that a commercially available RT-qPCR kit shows significant variabilities in expression profiles of the six salivary miRNA targets among the ten healthy participants. However, we demonstrate that the majority of these differences are associated to technical limitations rather than the biological contexts. Our data demonstrated that assessing the technical limitations of the kit coupled with a rigorous experimental design is crucial for interpreting biological relevance using RT-qPCR data. Regarding the high potential application of salivary miRNA in medical diagnosis, further technological breakthroughs for reliable tests are still required to overcome data inconsistencies and study dependent results.

## Aknowledgements

The authors would like to thank all participants who made this study possible. We sincerely thanks Victor Petit, CEO of the company SkillCell for all the insightful discussion and comments from the beginning till the manuscript writing process. We thanks the CNRS for being the promotor and the financial supporter of the study.

## Author contributions

Study concept and design: F.M. and TNN.V. Acquisition, analysis, interpretation of data: All authors. Writing of the manuscript and critical revision for important intellectual content: M.VDH., F.M. and TNN.V. Statistical analysis: M.VDH. Funding acquisition: F.M. All authors critically reviewed and edited the manuscript.

## Data availability statement

The authors declare that all relevant data has been provided within the manuscript and its supporting information files.

