## Supplementary Information for "Assessment of salivary microRNA by RT-qPCR: Challenges in data interpretation for clinical diagnosis"

Marc Van Der Hofstadt[1], Anna Cardinal[1], Morgane Lepeltier[1], Jérémy Boulestreau[1], Alimata Ouedraogo[1], Malik Kahli[1], Pierre Champigneux[1], Laurence Molina[1], Franck Molina[1*], Thi Nhu Ngoc Van[1*].

[1]Sys2Diag UMR9005 CNRS/ALCEN, Cap Gamma, Parc Euromédecine, 1682 rue de la Valsière, 34184, Montpellier, CEDEX 4, France.

**Keywords:** Saliva, miRNA, Biomarkers, Diagnostic, RT-qPCR.

**Table S1. The miRNAs chosen in this study.** Two similar pairs of miRNAs were included in this study, where they only differentiated by a single nucleotide (red font for the first pair and blue for the second pair).

| No | Accession miRBase | Name of mature miRNA | Sequence |
|---|---|---|---|
| 1 | MIMAT0000062 | hsa-let-7a-5p | UGAGGUAGUAGGUUGUAUAGUU |
| 2 | MIMAT0000067 | hsa-let-7f-5p | UGAGGUAGUAGAUUGUAUAGUU |
| 3 | MIMAT0000243 | has-miR-148a-3p | UCAGUGCACUACAGAACUUUGU |
| 4 | MIMAT0000083 | has-miR26b-5p | UUCAAGUAAUUCAGGAUAGGU |
| 5 | MIMAT0000104 | has-miR-107 | AGCAGCAUUGUACAGGGCUAUCA |
| 6 | MIMAT0000101 | has-miR-103a-3p | AGCAGCAUUGUACAGGGCUAUGA |

**Table S2. Average RT-qPCR Ct values of the four sampling points for each miRNAs assays assessed on the 10 participants.**

| | miRNA assay | | | | | |
|---|---|---|---|---|---|---|
| Participant | hsa-let-7a-5p | hsa-let-7f-5p | has-miR-148a-3p | has-miR26b-5p | has-miR-107 | has-miR-103a-3p |
| P1 | 24.79 ± 0.79 | 25.37 ±1.05 | 25.50 ± 1.10 | 24.10 ± 1.47 | 25.65 ± 0.90 | 25.22 ± 0.83 |
| P2 | 23.62 ± 1.37 | 23.94 ± 1.18 | 24.34 ± 1.28 | 22.16 ± 0.92 | 24.30 ± 1.65 | 23.78 ± 0.52 |
| P3 | 24.10 ± 1.75 | 24.54 ± 1.11 | 25.38 ± 0.84 | 25.07 ± 0.83 | 25.81 ± 0.91 | 25.19 ± 0.93 |
| P4 | 24.37 ± 1.60 | 24.53 ± 1.15 | 25.81 ± 0.80 | 24.38 ± 0.80 | 25.45 ± 1.17 | 24.96 ± 1.02 |
| P5 | 26.32 ± 0.76 | 26.97 ± 0.73 | 27.40 ± 1.05 | 27.46 ± 0.68 | 27.98 ± 1.21 | 27.37 ± 0.59 |
| P6 | 25.55 ± 0.71 | 27.09 ± 0.77 | 27.35 ± 0.76 | 27.37 ± 0.79 | 28.26 ± 1.03 | 27.52 ± 0.46 |
| P7 | 24.38 ± 0.63 | 24.77 ± 1.28 | 26.33 ± 0.93 | 25.51 ± 1.20 | 26.77 ± 1.99 | 25.70 ± 1.39 |
| P8 | 25.69 ± 0.11 | 24.25 ± 0.61 | 26.62 ± 0.50 | 25.95 ± 0.61 | 27.12 ± 0.60 | 26.67 ± 0.95 |
| P9 | 25.88 ± 1.10 | 25.04 ± 1.53 | 27.09 ± 1.23 | 25.67 ± 1.31 | 27.07 ± 1.61 | 26.71 ± 1.36 |
| P10 | 25.06 ± 0.45 | 24.99 ± 0.98 | 25.76 ± 0.86 | 24.61 ± 1.11 | 26.27 ± 0.96 | 25.84 ± 1.41 |

**Table S3: Dilution factors required to achieve 50 ng for the RT-qPCR reaction for the 10 participants and their 4 sampling points.**

| Participant | 1st Sampling | 2nd Sampling | 3rd Sampling | 4nd Sampling | Average |
|---|---|---|---|---|---|
| P1 | 19.68 | 33.82 | 24.30 | 42.24 | 30.01 |
| P2 | 63.92 | 35.83 | 17.86 | 20.34 | 34.49 |
| P3 | 20.04 | 24.67 | 17.42 | 19.14 | 20.32 |
| P4 | 71.24 | 47.04 | 41.70 | 8.92 | 42.23 |
| P5 | 5.26 | 8.00 | 6.88 | 7.26 | 6.85 |
| P6 | 9.48 | 9.74 | 7.96 | 9.38 | 9.14 |
| P7 | 27.2 | 30.00 | 24.70 | 24.36 | 26.56 |
| P8 | 38.7 | 19.74 | 30.86 | 12.16 | 25.37 |
| P9 | 34.12 | 19.20 | 14.60 | 24.48 | 23.10 |
| P10 | 52.44 | 37.73 | 33.88 | 32.48 | 39.13 |

**Table S4: Limit of detection (LOD) and limit of quantification (LOQ) for the six miRNA assays used in this study.**

| miRNA | LOD (copies/µL) | LOQ (copies/µL) | Ct at LOQ |
|---|---|---|---|
| hsa-Let-7a-5p | 1 | $10^5$ | 24.16 |
| hsa-Let-7f-5p | 1 | $10^4$ | 26.89 |
| hsa-miR-148a-3p | 1 | $10^5$ | 24.83 |
| hsa-miR26b-5p | 1 | $10^5$ | 23.83 |
| hsa-miR-107 | 1 | $10^5$ | 22.59 |
| hsa-miR-103a-3p | 1 | $10^2$ | 32.08 |

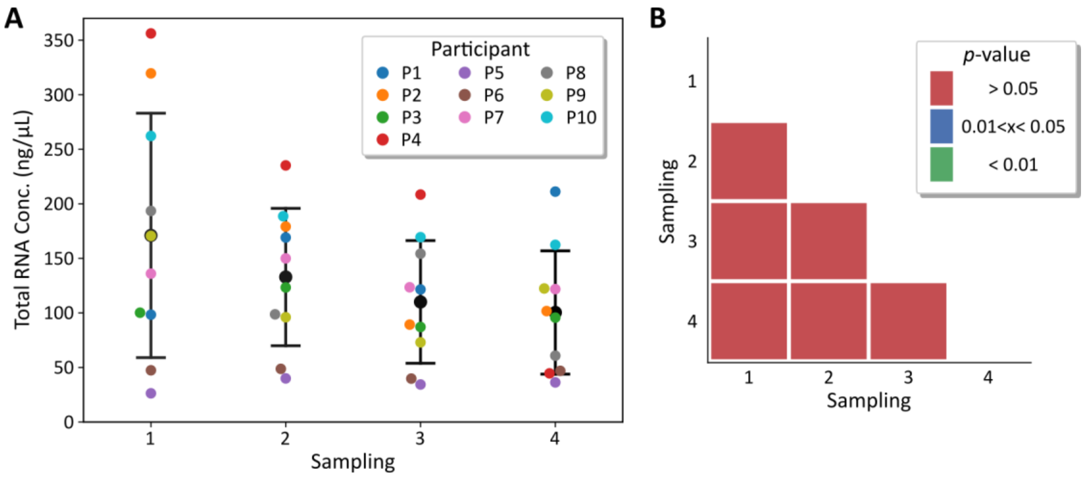


**Figure S1. Average total extracted small RNA concentration does not vary significantly among sampling points. (A)** Distribution of the total extracted small RNA concentrations among 10 participants throughout four sampling points. **(B)** Statistical analysis of panel A.

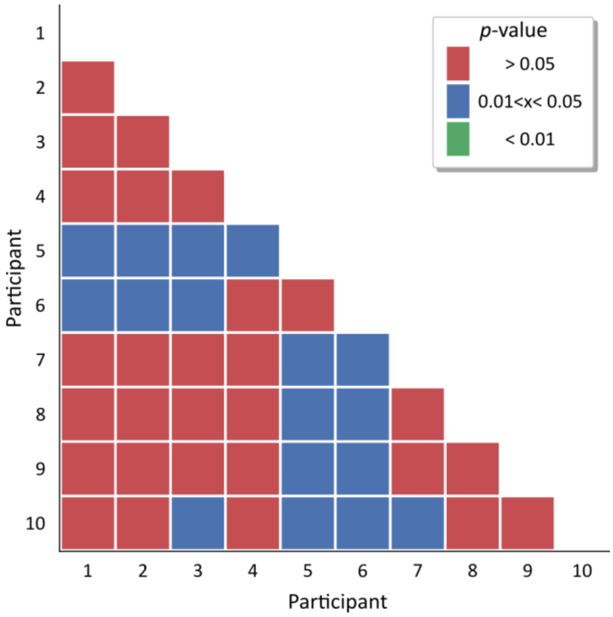


**Figure S2. Statistical analysis for the total extracted small RNA concentration highlighting the presence of a distinct group containing participant P5 and P6.**

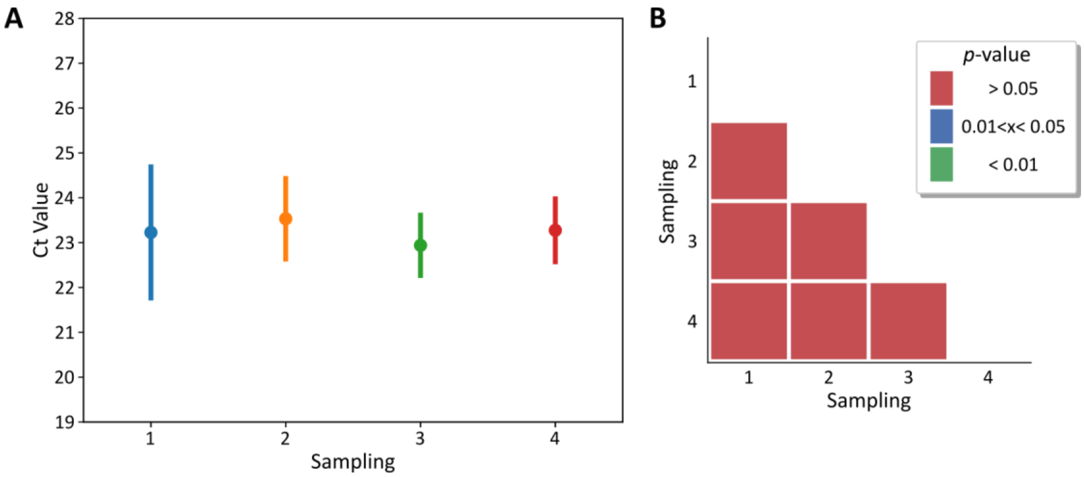


**Figure S3. Homogenous RNA extraction efficiencies throughout the study. (A)** RT-qPCR quantification values of spike-in UniSP6 miRNA using 50 ng of the total extracted salivary small RNAs for the different sampling points. **(B)** Statistical analysis for panel A.

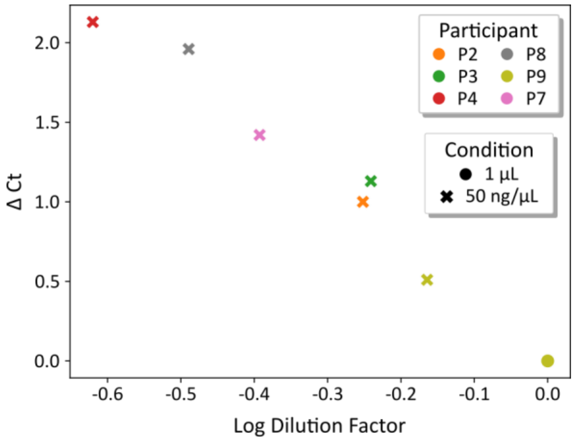


**Figure S4. Ct values of the spike-in UniSP6 miRNA quantified by RT-qPCR are associated with the dilution factors of the extracted small RNA samples.** All samples were analysed in two conditions: using 1µL (circle) or 50 ng (cross) of the total extracted small RNAs as input. Since the same concentration of UniSp6 miRNA is present at 1 µL for all participants, all 50 ng values have been normalized with respect to the RNA concentration at 1 µL (dilution factor), and all Ct values have been shifted with respect to the 1 µL value (causing the overlapping of all 1 µL values). ΔCt values = $Ct^{50ng} – Ct^{1\mu L}$.

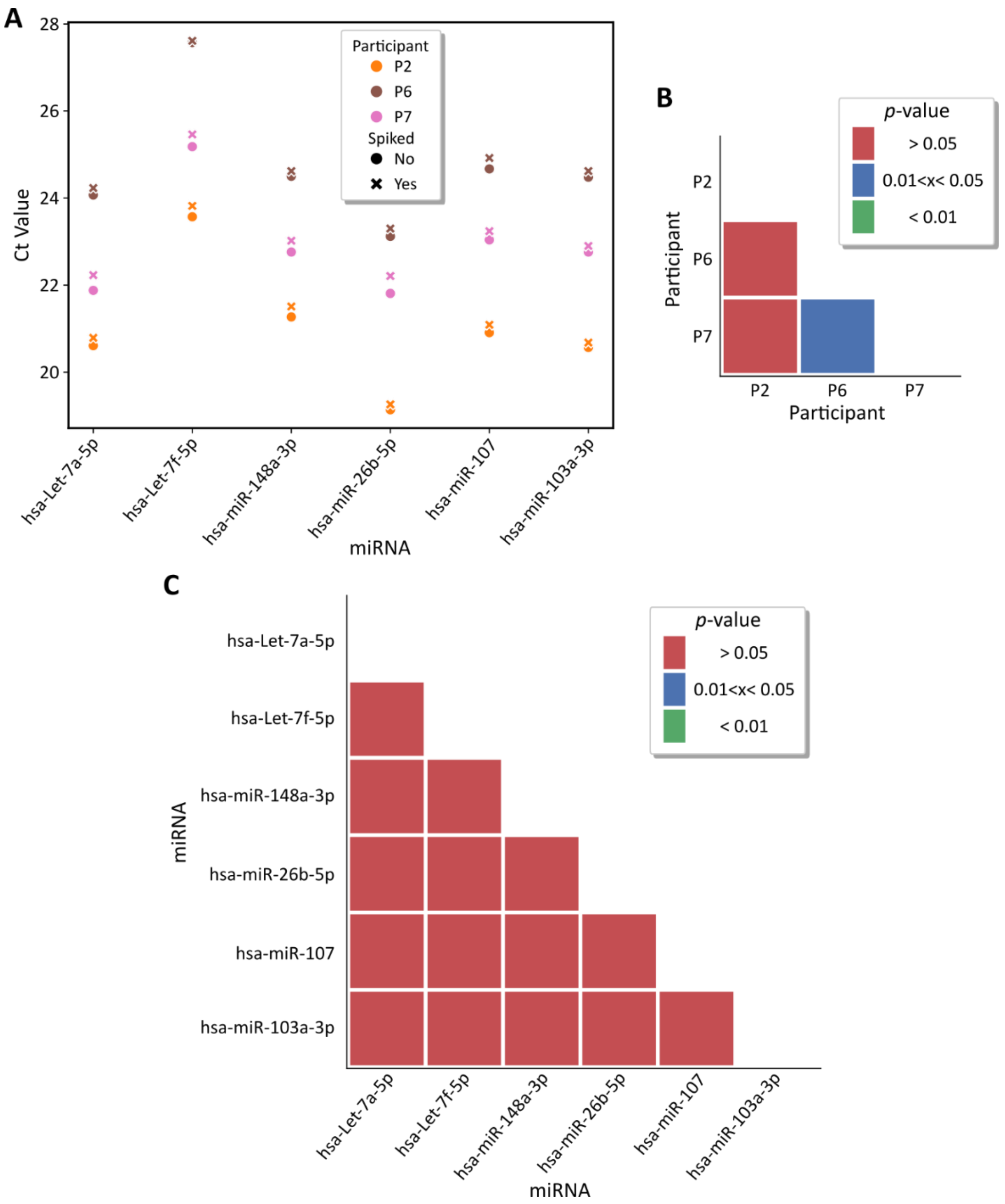


**Figure S5. Spike-in of artificial UniSP6 miRNA does not significantly interfere with neither six miRNA assays nor ten participants.** Small RNAs were extracted from three saliva samples (P2, P6 and P7) in the presence and absence of spiked-in artificial UniSP6 miRNA. (A) RT-qPCR quantification for the six miRNAs using 50 ng of small RNAs. Statistical analysis for panel A of ΔCt between with and without spiking with respect to (B) individual participants and (C) miRNA assay.

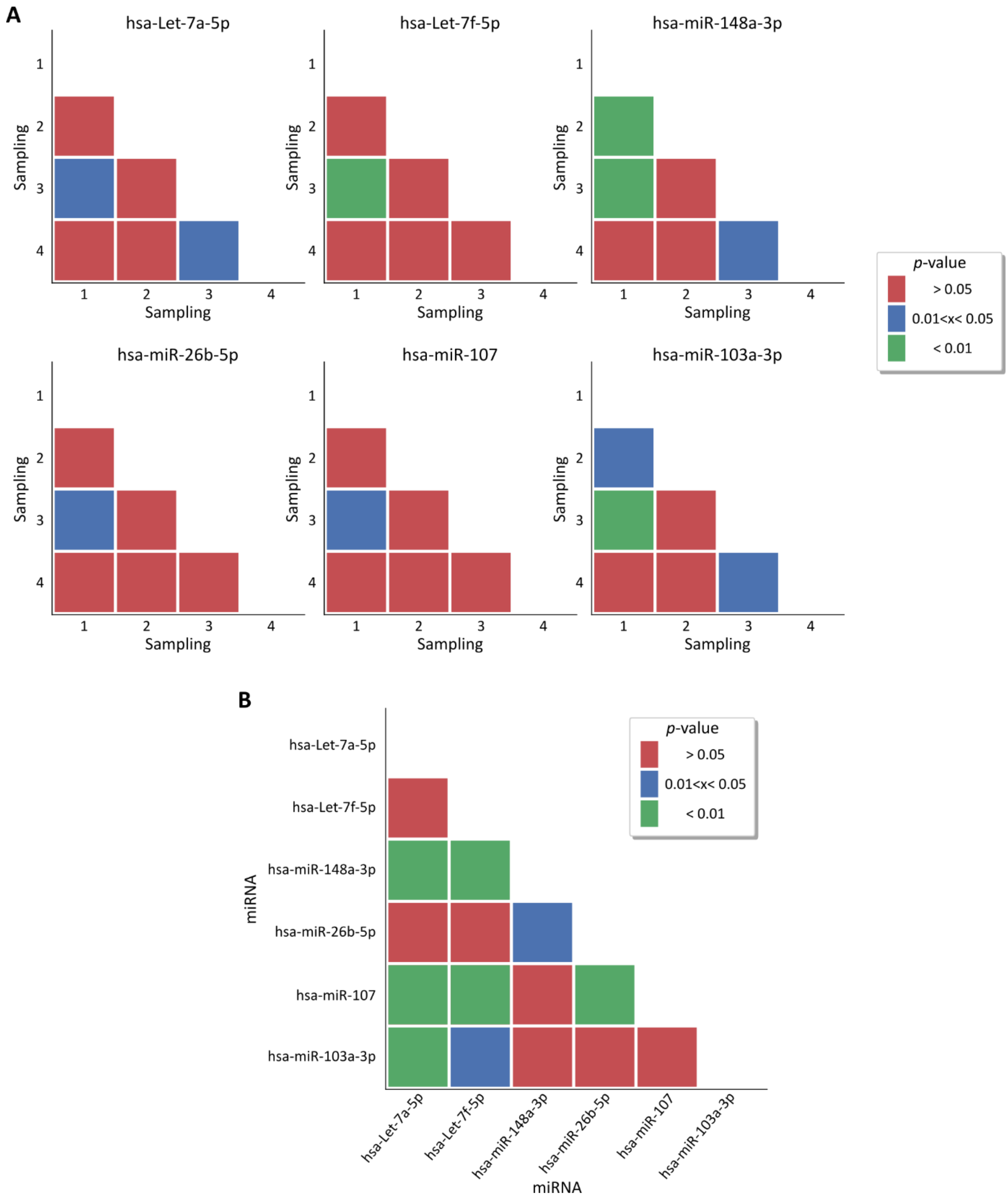


**Figure S6. Average Ct values analysing 50 ng small RNA input were dependent on the sampling point and the miRNA assay used.** Statistical analysis for data presented in Figure 2B of the manuscript showing differences within **(A)** samplings for each miRNA assay and **(B)** within miRNA assays.

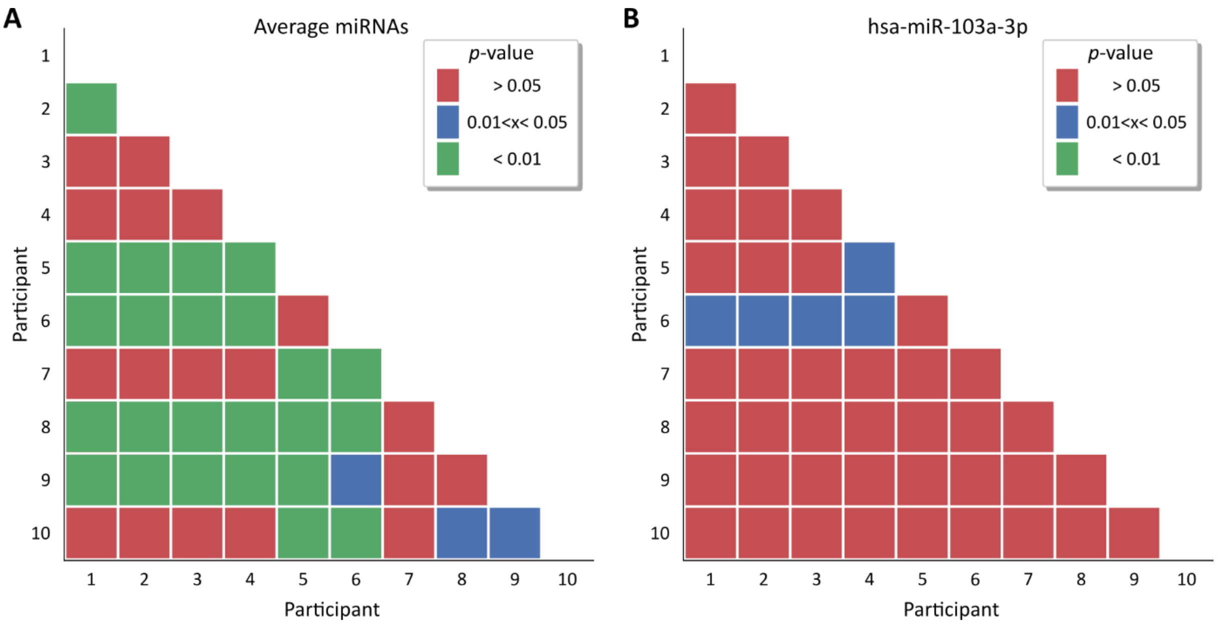


**Figure S7. Statistical analysis for the Ct values (Figure 2C of the manuscript) highlighting the division of the participants upon miRNA expression.** Statistical analysis for data in Figure 2C showing statistical differences within participants **(A)** when averaging the six miRNAs and **(B)** within hsa-miR-103a-3p. We note that in panel A participant P7 and participant P10 do not clearly belong to a group but rather are adjacent to a group, sharing partial significance.

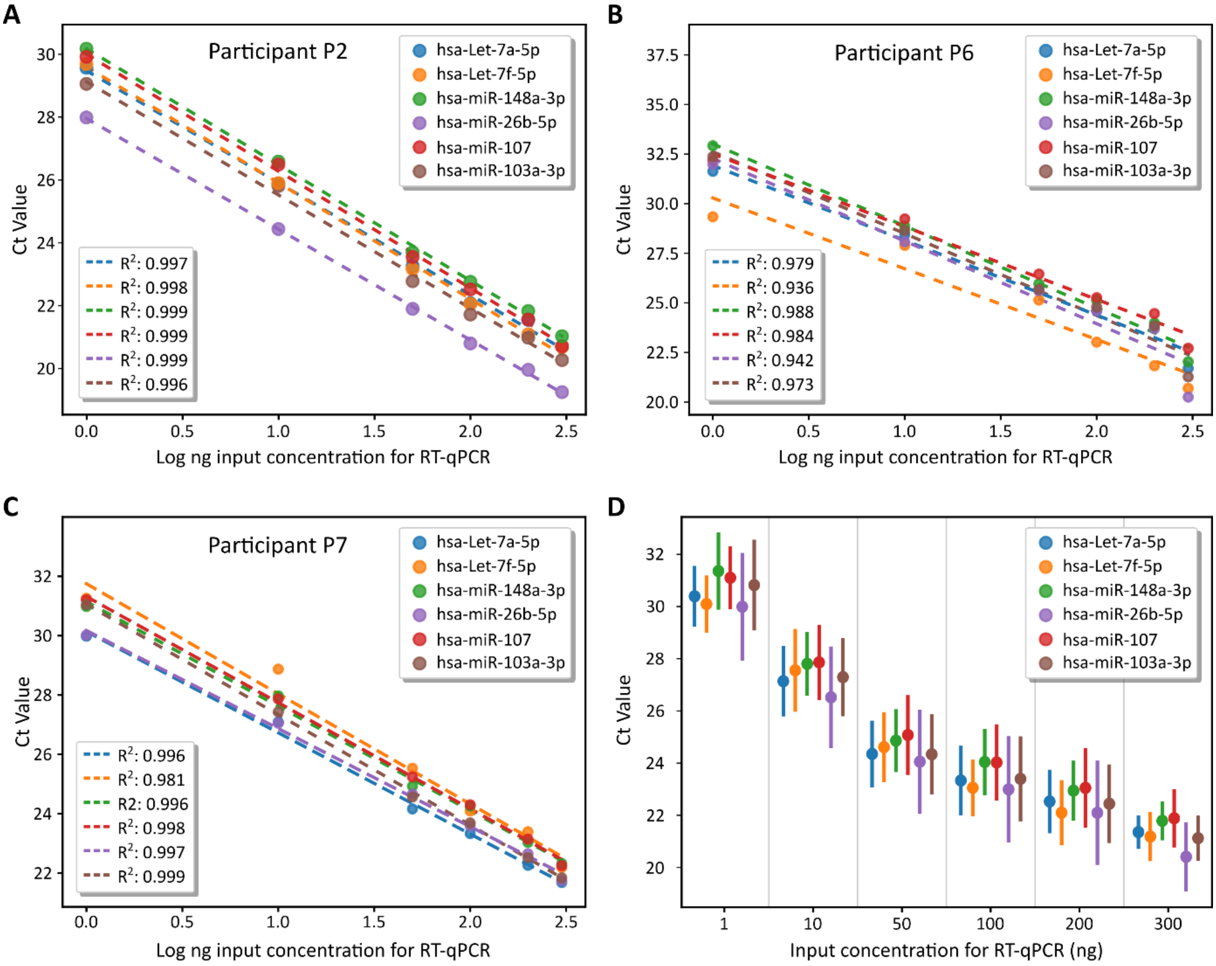


**Figure S8. Dose dependent responses of the six miRNAs assays from three participants P2 (A), P6 (B) and P7 (C).** Average response in Ct values was calculated **(D)**. 1 ng, 10 ng, 100 ng, 200 ng and 300 ng of total extracted small RNA from each participant were analysed by the six miRNA assays.

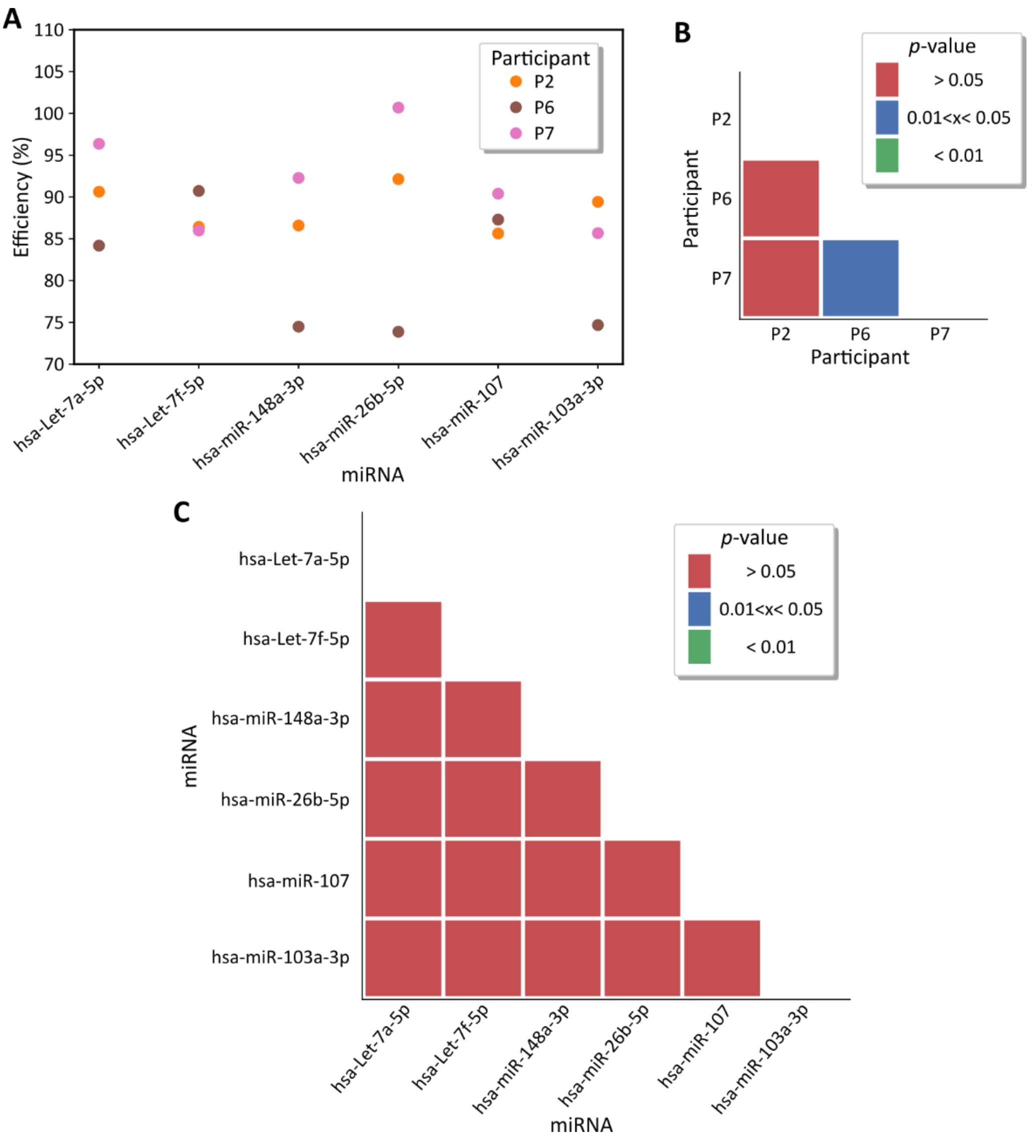


**Figure S9. Homogeneity in RT-PCR efficiency of miRNA assays for participants P2 and P7 contrary to participant P6 analysing extracted small RNA from saliva samples of these participants. (A)** RT-PCR efficiencies calculated from the dose dependent data presented in Figure S5. Statistical analysis for panel A with respect to **(B)** participants and **(C)** miRNA assay.

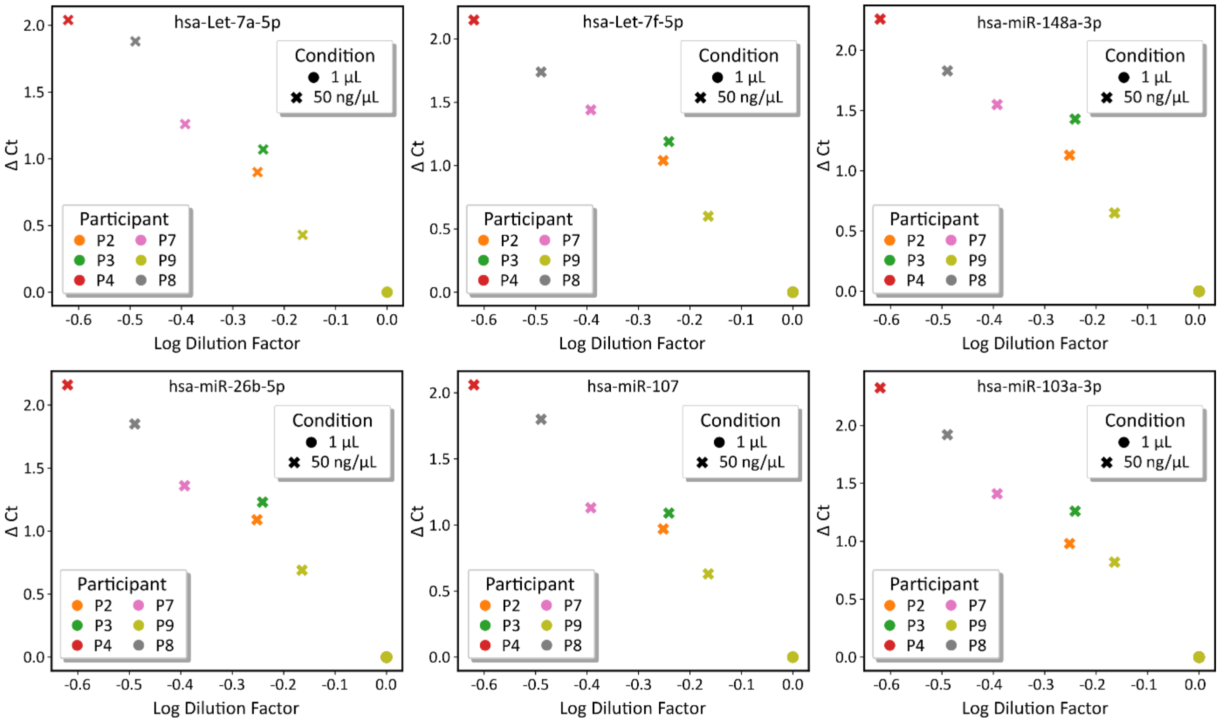


**Figure S10. The dilution factor required to achieve 50 ng of small RNA input does not affect RT-qPCR efficiency of the six miRNA assays.** As for figure S4, two inputs concentrations where used, 1µL (circle) or 50 ng (cross). Again, all 50 ng values have been normalized with respect to the RNA concentration at 1 µL (dilution factor) and all Ct values have been shifted with respect to the 1 µL value. Although in this case the miRNA concentration may or not be heterogeneous, a linear behaviour is still obtained because we are observing the linear behaviour on the Ct value (ΔCt) due to the dilution effect (log dilution factor). ΔCt values = $Ct^{50ng} – Ct^{1\mu L}$.

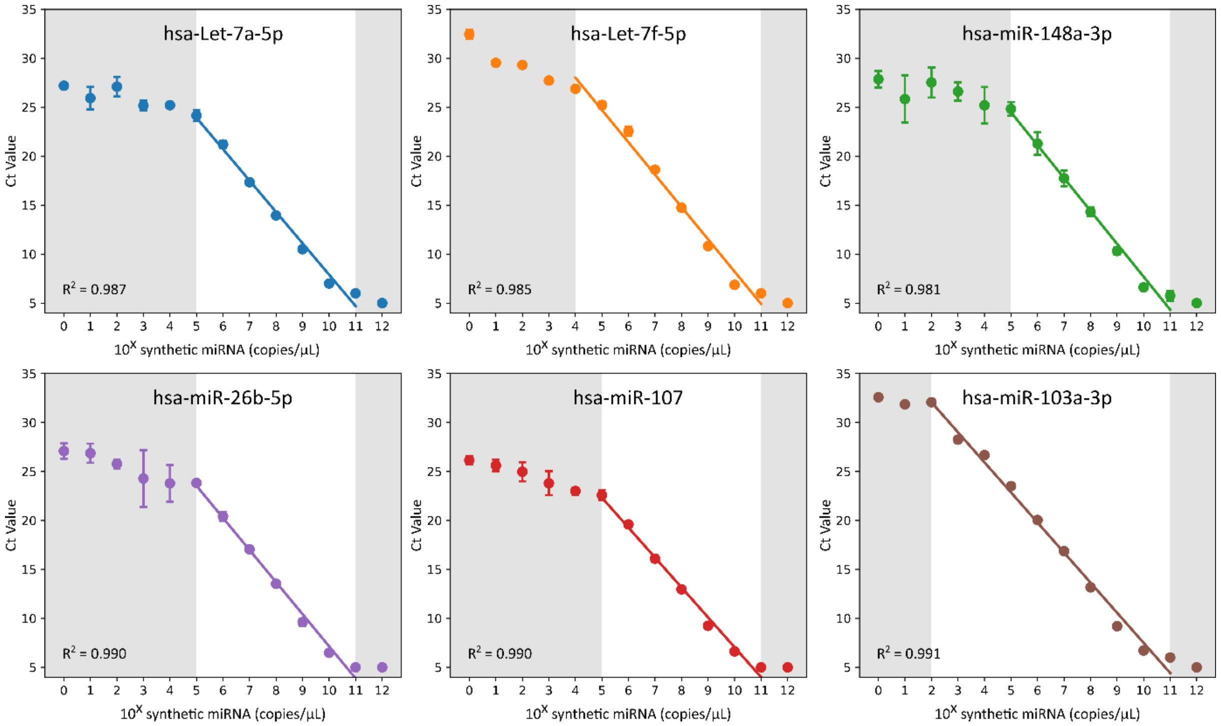


**Figure S11. The limit of detection (LOD) and quantification (LOQ) differ for the six miRNA RT-qPCR assays.** Individual plots for data shown in Figure 3A of the manuscript, demonstrating high sensitivity (LOD of 1 copy/µL) for all six assays. Regression line calculated within LOQ and Ct saturation point (*i.e.* Ct value = 5). The grey region delimits the quantification region.

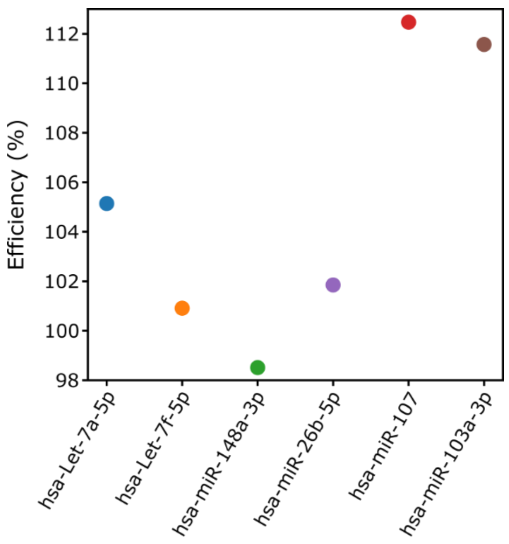


**Figure S12. RT-qPCR efficiency for the six miRNA assays analysing their synthetic targets.** Data from Figure S11 was used to calculate efficiencies.

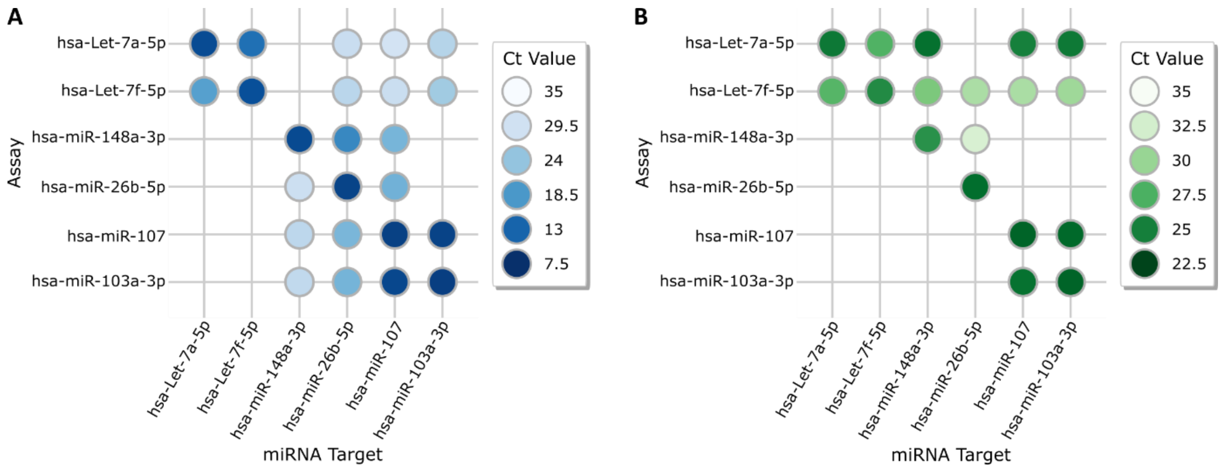


**Figure S13. High cross-detection among the six miRNA assays used in this study**. Cross detections among miRNA assays when using **(A)** $10^9$ copies/µL and **(B)** $10^5$ copies/µL synthetic targets. The absence of circles indicate no detection or Ct values = 35. This data was used to calculate the ΔCt values in Figure 4B & 4C of the manuscript.

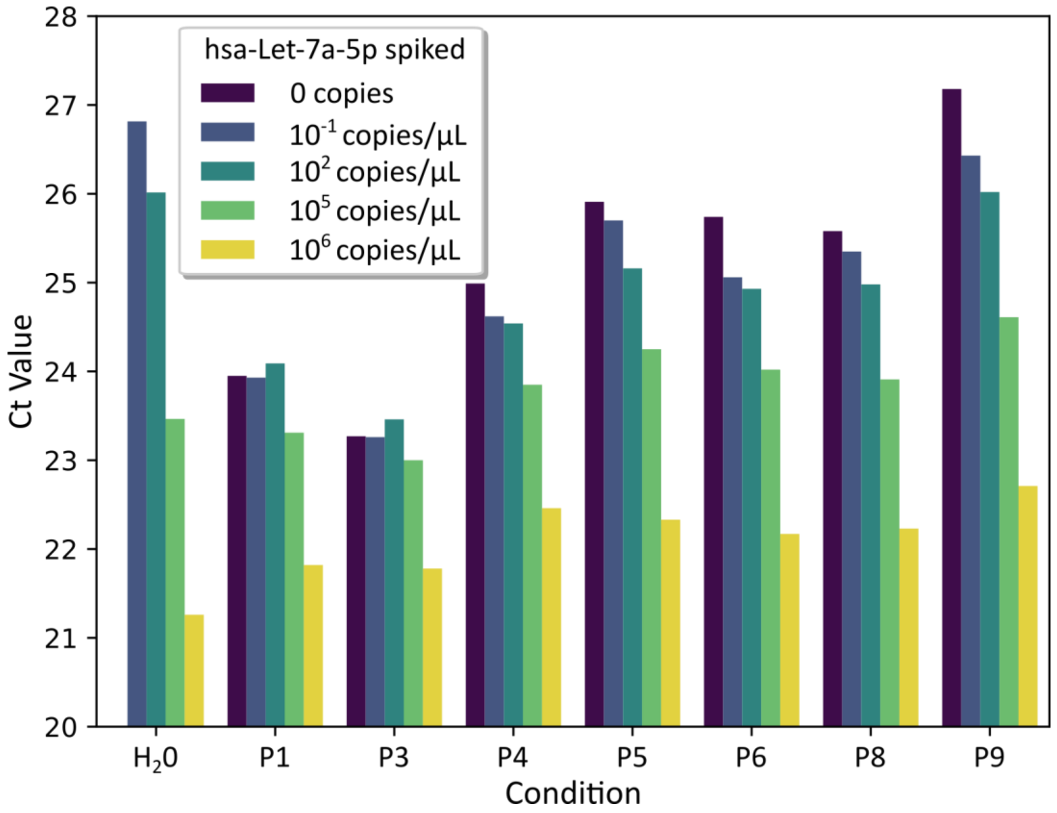


**Figure S14. The spike-in of synthetic hsa-lest-7a-5p miRNA decreases the Ct Values of extracted small RNA samples**. Data used to calculate the ΔCt values in Figure 5 of the manuscript.
